# Horizontally transmitted symbiont populations in deep-sea mussels are genetically isolated

**DOI:** 10.1101/536854

**Authors:** Devani Romero Picazo, Tal Dagan, Rebecca Ansorge, Jillian M. Petersen, Nicole Dubilier, Anne Kupczok

## Abstract

Eukaryotes are habitats for bacterial organisms where the host colonization and dispersal among individual hosts have consequences for the bacterial ecology and evolution. Vertical symbiont transmission leads to geographic isolation of the microbial population and consequently to genetic isolation of microbiotas from individual hosts. In contrast, the extent of geographic and genetic isolation of horizontally transmitted microbiota is poorly characterized. Here we show that chemosynthetic symbionts of individual *Bathymodiolus brooksi* mussels constitute genetically isolated populations. The reconstruction of core genome-wide strain sequences from high-resolution metagenomes revealed distinct phylogenetic clades. Nucleotide diversity and strain composition vary along the mussel lifespan and individual hosts show a high degree of genetic isolation. Our results suggest that the uptake of environmental bacteria is a restricted process in *B. brooksi*, where self-infection of the gill tissue results in serial founder effects during symbiont evolution. We conclude that bacterial colonization dynamics over the host life-cycle is thus an important determinant of population structure and genome evolution of horizontally transmitted symbionts.

---

Bacteria inhabit most eukaryotes where their presence has consequences for key aspects of their host biology^1^, such as host development^2^, nutrition^3^, or behavior^4^. From the bacterial perspective, animals constitute an ecological niche, where microbial communities utilize the resources of their host habitat^5^. The microbiota biodiversity over the host life cycle is determined by bacteria colonization dynamics and by host properties, including biotic and abiotic factors. For example, the microbiota can be affected by the host diet^6^ or the host physiological state (e.g., hibernation^7^ or pregnancy^8^). In addition, changes in the host environmental conditions such as temperature^9^ or the availability of reduced compounds^10^ can have an effect on the microbiota community composition.

Microbiota dispersal over the host life cycle depends on the level of fidelity between the host and its microbiota; in faithful interactions, vertically transmitted bacteria are transferred from adults to their progeny during early host developmental stages, while in less faithful interactions, horizontally transmitted bacteria are acquired from the environment throughout the host life cycle^11^. Strictly vertically transmitted bacteria are specialized in their host niche and their association with the host imposes an extreme geographic isolation. Bacterial inheritance over host generations imposes a strong bottleneck on the microbiota population and leads to reduced intra-host genetic diversity^12^. Examples are monoclonal or biclonal populations observed in symbiotic bacteria inhabiting grass sharpshooter^13^ and pea aphids^14^. Furthermore, the geographic isolation of vertically transmitted bacteria leads to genetic isolation and to symbiont genome reduction over time as a consequence of genetic drift^15^. In contrast, dispersal is expected to be higher for horizontally transmitted bacteria, where host-associated populations are connected to one another through the environmental pool^16^. Nonetheless, the genetic diversity of horizontally transmitted microbial populations may also be reduced due to bottlenecks during symbiont transmission and host colonization. Stochastic effects in the colonization of horizontally transmitted bacteria may manifest themselves in differences in microbiota strain composition among hosts^17,18^. This would lead to subdivided symbiont populations where the geographic isolation of the microbiota depends on the degree of symbiont dispersal among individual hosts. Geographic isolation of individual hosts over the host life span would then lead to genetic isolation of the symbiont populations and to symbiont population structure. Genomic variation and genetic isolation have been observed for horizontally transmitted symbionts of the human gut microbiome^19^ and of the honey bee gut microbiome^20^. Moreover, structured symbiont populations can also emerge within an individual host, as observed for *Vibrio fischeri* colonizing the squid light organ, where different light organ crypts are infected by a specific strain^21^. The degree of dispersal of horizontally transmitted symbionts remains understudied; hence, whether populations from different microbiomes are intermixing or are genetically isolated is generally unknown.

Here we study the microbiota strain composition of horizontally transmitted endosymbionts across individual *Bathymodiolus brooksi* deep-sea mussels. *Bathymodiolus* mussels live in a nutritional symbiosis with the chemosynthetic sulfur-oxidizing (SOX) and methane-oxidizing (MOX) bacteria. The symbionts are acquired horizontally from the seawater and are harbored in bacteriocytes within the gill epithelium^22,23^. Most Bathymodiolus species harbor only a single 16S rRNA phylotype for each symbiont, including *B. brooksi*^24^. Nevertheless, a recent metagenomic analysis of *Bathymodiolus* species from hydrothermal vents in the mid Atlantic ridge showed the presence of different SOX strains with differing metabolic capacity^25^. Mussel gills constantly develop new filaments that are continuously infected^26^. However, whether the new gill filaments in *Bathymodiolus brooksi* are colonized predominantly by environmental bacteria or by symbionts from older filaments of the same host remains unknown. These two alternative scenarios are expected to impose different degrees of geographic isolation on the symbiont population: in continuous environmental acquisition, the level of inter-host dispersal is high while self-infection limits the symbiont dispersal. Here we studied the impact of tissue colonization dynamics of horizontally transmitted intracellular symbionts on the degree of symbiont diversity. Furthermore, we quantified the level of genetic isolation among communities across individual mussels and its impact on symbiont genome evolution. For that, we implemented a high-resolution metagenomics approach that captures genome-wide diversity for both symbionts in multiple *Bathymodiolus brooksi* individuals from a single site.

## Results

### Gene-based metagenomics binning recovers SOX and MOX core genomes

To study the evolution of the SOX and MOX genomes in *Bathymodiolus* mussels we used a high-resolution metagenomics approach. Twenty-three *B. brooksi* individuals of shell sizes ranging between 4.8 cm and 24.3 cm were sampled from a single location at a cold seep site in the northern Gulf of Mexico. Shell size correlates with mussel age^27^; thus, analyzing mussels within a wide shell size range allowed us to study the symbiont population structure across host ages. The mussels were sampled from three separate mussel ‘clumps’ (small mussel patches residing on the sediment) that were at most 131m apart (Supplementary Fig. 1). Such a ‘patchy’ distribution has often been observed in deep-sea mussels^28^. To obtain a comprehensive representation of the bacterial population in individual mussels and to accurately infer strain-specific genomes, homogenized gill tissue of each mussel was deeply sequenced (on average, 37.8 million paired-end reads of 250bp per sample, Supplementary Table 1). The resulting metagenomic sequencing data was analyzed by a gene-based binning approach^29^.

The prediction of protein-coding genes from the assembled metagenomes yielded a non-redundant gene catalog of 4.4 million genes that potentially contains every gene present in the samples. This includes genes from the microbial community and from the mussel host. In the metagenomics binning step, genes that covary in their abundance across the different samples were clustered into metagenomic species (MGSs). Our analysis revealed two MGSs that comprise the SOX and MOX core genomes (Supplementary Fig. 2). The distribution of gene coverage in individual samples shows that genes in each core genome have a similar abundance within each mussel. This confirms the classification of the SOX and MOX MGSs as core genomes. The MOX core genome is the largest MGS and it contains 2,518 genes with a total length of 1.97 Mbp. A comparison to Gammaproteobacteria marker genes shows that it is 96.2% complete. Furthermore, it contains 1,568 genes (62.3%) that have homologs in MOX-related genomes. The SOX core genome contains 1,439 genes, has a total length of 1.27 Mbp and is considered as 80.2% complete. It contains 1,188 genes (82.6%) with homologs in SOX-related genomes. In addition to the SOX and MOX core genomes, our analysis revealed a third MGS of 1,449 genes (Supplementary Fig. 2) that was found in low abundance in a single mussel and, in addition, 98,944 co-abundant gene groups (CAGs, 3-699 genes). Of the 23 metagenomes, four samples were discarded during the metagenomics binning. Two samples were discarded prior to the binning due to high variance in symbiont marker gene coverages and two samples were discarded after binning due to low coverage for both symbionts (Supplementary Figs. 2,3). To gain insight into the SOX and MOX population structure between hosts, we compared the characteristics of the core genomes across the remaining 19 samples. The analysis of the core genome coverages shows that SOX is the dominant member of the mussel microbiota. The differences in the SOX to MOX ratio among the mussel metagenomes are likely explained by differences in the availability of H_2_S and CH_4_ among clumps, which is a known determinant of SOX and MOX abundance in *Bathymodiolus*^30^ (Supplementary Information, Supplementary Fig. 4).

To study symbiont diversity below the species level, we analyzed single nucleotide variants (SNVs) that were detected in the core genomes of the two symbionts. In this analysis, we considered SNVs that are fixed in a metagenome as well as polymorphic SNVs, i.e., SNVs, where both the reference and the alternative allele are observed in a single metagenome. We found 18,070 SNVs in SOX (SNV density of 14 SNVs/kbp, 49 multi-state, 0.27%) and 4,652 SNVs in MOX (SNV density of 2.4 SNVs/kbp, 5 multi-state, 0.11%). The number of polymorphic SNVs per sample ranges from 162 (0.9%) to 11,064 (61%) for SOX and from 27 (0.58%) to 3,026 (65%) for MOX (Supplementary Table 1), thus, most SNVs are polymorphic in at least one sample. It is important to note that the observed difference in strain-level diversity between SOX and MOX cannot be explained by the difference in sequencing depth (Supplementary Information). These results are in agreement with previous reports of SOX genetic diversity in other *Bathymodiolus* species^25^. We further revealed that there is genetic diversity in the MOX symbiont.

### Bathymodiolus microbiota is composed of SOX and MOX strains from several clades

Diversity in natural populations of bacteria is characterized by cohesive associations among genetic loci that contribute to lineage formation and generate distinguishable genetic clusters beyond the species level^31^. The formation of niche-specific genotypes (i.e., ecotypes) has been mainly studied in populations of free-living organisms such as the cyanobacterium *Prochlorococcus* spp.^32^. Here we consider a strain to be a genetic entity that is present in multiple hosts and is characterized by a set of clustered variants in the core genome. To study lineage formation in symbiont populations associated with *Bathymodiolus* mussels, we reconstructed the strain consensus core genomes from strain-specific variants that show similar frequencies in a metagenomic sample.

The SNVs found in multiple samples and their covariation across samples were used for strain deconvolution of the core genomes using DESMAN^33^. This revealed that SOX is composed of eleven different strains with a mean strain core genome sequence identity of 99.52%. Phylogenetic reconstruction shows that the eleven strains cluster into four clades, which are separated by relatively long internal branches (Fig. 1b). Notably, 849 of the SNVs on the SOX core genome (4.7%) do not differentiate between strains. Thus, the resulting strain alignment is invariant for each of these positions and they are termed invariant SNVs from here on. For MOX, six strains with a mean core genome sequence identity of 99.88% were reconstructed. The phylogenetic network shows that the six strains cluster into two clades comprising three strains each (Fig. 1e). Of the total SNVs, 1,138 (24.4%) are invariant in the strain alignment. The overall MOX branch lengths are shorter than those of SOX. We detected no effect of sequencing coverage on the inference of the strain clades for SOX (Supplementary Information, Supplementary Fig. 5).

**Figure 1:**
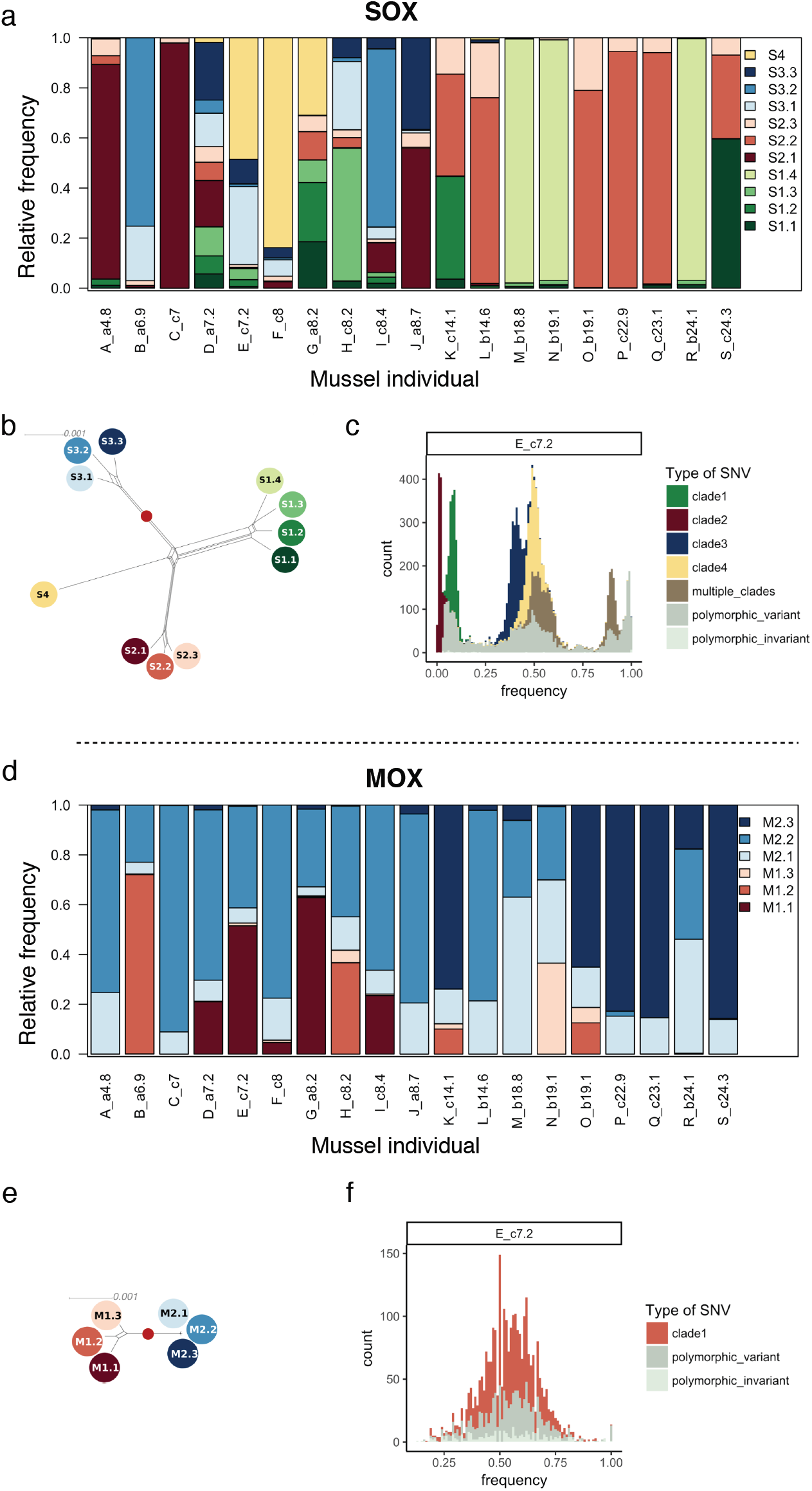
Symbiont strain abundances (a, d), symbiont strain relationships (b, e) and example allele frequency spectra (c, f). **a**, **b**, **c**, 11 strains reconstructed for SOX. These cluster into four clades, with two times four, three and one strain per clade, labelled by shades of green, red, blue, and yellow. The strains differ by between 669 SNVs (strains S2.2 and S2.3, sequence identity 99.95%) and 8,171 SNVs (strains S3.2 and S4 sequence identity 99.36%). Minimum number of SNVs between strains of different clades is 6,451 (strains S1.1 and S2.1, sequence identify 99.49%). **d**, **e**, **f**, 6 strains reconstructed for MOX. These cluster into two clades, labelled by shades of red and blue. Strains differ by between 105 (strain M2.2 and M2.3, sequence identity 99.99%) and 2,677 SNVs (strain M1.1 and M2.1, sequence identity 99.81). The minimum number of SNVs differentiating strains from different clades is 2,224 (strains M2.2 and M1.3, sequence identity 99.85%). **a**, **d**, Stacked barplot of strain relative abundances for each individual mussel. Mussel individuals are labeled with an assigned letter (A-S), followed by the sampling clump (a, b or c) and the shell size (cm). **b**, **e**, Splits network of the strain genome sequences. Scale bar shows the number of differences per site. The red dots indicate the position of the root. **c**, **f**, Example of derived allele frequency spectra (sample E). Different colors represent different strain clades (see also Supplementary Fig. 6).

To study the community assembly at the strain level, we examined the strain distribution across individual mussels. Each SOX strain could be identified in between three and eight samples (frequency ≥5%; Fig. 1a). Only one or two strains were detected with a frequency of at least 5% in small mussels (≤7 cm), two to nine strains in medium-sized mussels (7.2 cm – 14.1 cm) and one to two strains in large mussels (14.6 cm – 24.1 cm). Notably, only strains from clades S1 and S2 are present in large mussels (≥14.6 cm). One of the large mussels (S) is an exception as it hosts three SOX strains and contains strains from both clades S1 and S2. Six mussels have one dominant SOX strain (frequency ≥90%). Five of these are large mussels (M, N, P, Q, R) and only one is a small mussel (C). The dominant strain is either S1.4, S2.1, or S2.2 (Fig. 1a; Supplementary Table 1). The MOX strain composition across mussels shows that each MOX strain occurs (frequency ≥5%; Fig. 1d) in four to 17 mussels and each mussel contains two to four MOX strains. Additionally, strains of clade M2 are dominant in ten of the mussels.

To investigate the degree of genetic cohesion within strain clades in the population, we studied the allele frequency spectrum (AFS) of each mussel. A visual inspection of the derived allele frequency spectra revealed multimodal distributions for both symbiont populations. The modes reach high allele frequencies and are associated with the main phylogenetic clades; this suggests that the clades constitute cohesive genetic units (Fig. 1c,f; Supplementary Fig. 6). The presence of high-frequency modes is especially apparent for SOX in medium-sized mussels that contain multiple strains. To identify sample-specific strain sequences, we reconstructed dominant haplotypes (major allele frequency ≥90%) for the samples that contain a dominant strain (strain frequency ≥90%). By comparing dominant haplotypes among samples containing the same dominant strain, we found that these can contain between 42 and 74 differential SNVs (Supplementary Table 1). This suggests that the fixation of variants within individual mussels contributes to the observed population structure.

Overall, our results revealed that the symbiont populations are composed of strains that cluster into few clades, which appear to be maintained by strong cohesive forces. In addition, the strains are shared among multiple mussels and multiple strains are capable of dominating different hosts. This suggests that stochastic processes are governing the symbiont community assembly, as previously proposed for other *Bathymodiolus* species^34^.

### SOX strains evolve under purifying selection while MOX evolution is characterized by neutral processes

To study the evolution of SOX and MOX strains in *Bathymodiolus*, we examined the selection regimes that have been involved in the formation of cohesive genetic SOX and MOX units. The core genome-wide ratio of pN/pS is higher in MOX (pN/pS of 0.425) in comparison to SOX (pN/pS of 0.137), which indicates that the strength of purifying selection is higher for SOX. In addition, we estimated pN/pS for each of the symbiont core genes. This revealed that MOX genes are characterized by large pN/pS and small pS values, while SOX genes have small pN/pS and large pS values (Supplementary Fig. 7). The relative rate of nonsynonymous to synonymous substitutions has been shown to depend on the divergence of the analyzed species^35,36^. For populations of low divergence, SNVs comprise substitutions that have been fixed in the population and mutations that arose recently. The latter include slightly deleterious mutations that were not yet purged by selection, resulting in an elevated ratio of nonsynonymous to synonymous replacements. Thus, this ratio is not suitable for analyzing closely related genomes, which is usually the case when studying variation within bacterial species.

To circumvent the bias in pN/pS, we tested for differences in selection regimes in the evolution of SOX and MOX strains using the neutrality index (NI). NI is used to distinguish between divergent and polymorphic SNVs and to quantify the departure of a population from the neutral expectation. An excess of divergent nonsynonymous mutations (NI<1) indicates that the population underwent positive selection or an important demographic change in the past^37^. We estimated NI by considering two different levels of divergence and polymorphism. In the first level, all identified strains are considered as diverged taxonomic units; in the second level, we disregard the small-scale strain classification and consider only the clades as diverged taxonomic units (Table 1). Considering all strains as divergent, we observed a low NI^MOX^ (<1), which suggests that MOX evolved under a neutral (NI∼1) or positive selection regime. NI^MOX^ increased when considering the clades as diverged, which suggests that the low NI^MOX^ observed at the strain level is the result of an excess of nonsynonymous SNVs within the strain clades that may constitute transient polymorphisms. Thus, the excess of nonsynonymous mutations observed for MOX is biased by the low level of divergence; hence, similar to the pN/pS ratio, it cannot serve as an indication for positive selection. On the other hand, we found that purifying selection is in action for SOX (NI^SOX^>1). Similar to MOX, when using the clades as divergent, NI^SOX^ slightly increases. This indicates that the SNVs that differ between clades are more likely to be substitutions in comparison to those that differ among within-clade strains.

**Table 1.** Neutrality index (NI) for the symbiont core genomes.

**a**
|  | SOX |  | MOX |  |
| --- | --- | --- | --- | --- |
|  | divergent | polymorphic | divergent | polymorphic |
| nonsynonymous SNVs | 5004 | 990 | 2115 | 704 |
| synonymous SNVs | 10577 | 1450 | 1313 | 515 |
| nonsynonymous SNVs/synonymous SNVs | 0.47 | 0.68 | 1.61 | 1.37 |
| NI | 1.44 |  | 0.85 |  |

**b**
|  | SOX |  | MOX |  |
| --- | --- | --- | --- | --- |
|  | divergent | polymorphic | divergent | polymorphic |
| nonsynonymous SNVs | 2549 | 3455 | 1041 | 1778 |
| synonymous SNVs | 6370 | 5657 | 649 | 1179 |
| nonsynonymous SNVs/synonymous SNVs | 0.40 | 0.61 | 1.60 | 1.51 |
| NI | 1.52 |  | 0.94 |  |

**a**, divergent SNVs are all those SNVs that differ between at least two strains, i.e., all identified strains are considered as diverged taxonomic units, and polymorphic SNVs are all the invariant SNVs. **b**, Divergent SNVs have the same state inside a strain clade and are not invariant and polymorphic SNVs are all the remaining, i.e., only the clades are considered as diverged taxonomic units.

Altogether, these results suggest differences in the selection regimes during the evolution of the SOX and MOX strains. While the SOX core genome is shaped by purifying selection, we cannot detect deviation from the neutral expectation in the MOX core genome. These differences likely stem from the different divergence levels among the strains of both symbiont populations. The association of SOX with *Bathymodiolus* mussels is considered to be ancient in chemosynthetic deep-sea mussels whereas the MOX association is thought to have evolved secondarily during *Bathymodiolus* diversification^38^. This is in agreement with the larger degree of divergence observed here for SOX. Since we observed no evidence for positive selection on the symbiont core genomes, we suggest that the strains constitute cohesive genetic units within one ecotype^39^, where all strains are functionally equivalent at the core genome level. Notwithstanding, the strains might be linked to differences in the accessory gene content, as observed, for example, in the free-living cyanobacterium *Prochlorococcus* spp.^32^ and in SOX symbionts of other *Bathymodiolus* species^25^.

### Intra-sample diversity is higher for SOX than for MOX

The association with the host limits the dispersal of bacterial populations where the association across generations is likely maintained by symbiont dispersal between host individuals. If symbionts are not continuously taken up from the environment, each individual host constitutes an isolated habitat over its lifetime^5^. Geographic isolation between habitats results in genetic isolation and contributes to the formation of cohesive associations of genetic loci^31^. Previous studies showed that geographic isolation during vertical transmission can lead to the reduction of intra-host genetic diversity in the bacterial populations^12^, nonetheless, the degree of isolation remains understudied for horizontally transmitted microbes. To characterize the contribution of geographic isolation to strain formation in the *Bathymodiolous* symbiosis, we next studied the degree of genetic isolation. Our sample collection of mussels covering a range of sizes (and thus ages) enabled us to compare symbiont genome diversity among individual hosts of different age within a single sampling site, thus minimizing the putative effect of biogeography on population structure. The host species *B. brooksi* is ideal for such an analysis as it grows to unusally large sizes and possibly lives longer than many other *Bathymodiolus* species. To study differences in genome diversity of the two symbionts across individual mussels, we estimated the intra-sample nucleotide diversity (π) and the ecological measure α-diversity at the resolution of the SOX and MOX strains.

We found a high variability of π^SOX^ among different mussels (intra-sample π^SOX^ between 5.2×10^-5^ and 3.6×10^-3^, Table 2, Fig. 2). Furthermore, π^SOX^ and the SOX α-diversity are significantly positively correlated (ρ^2^=0.98, p<10^-6^, Spearman correlation, Fig. 2a); hence, the intra-sample strain diversity is well explained by the nucleotide diversity. The variability in π^SOX^ agrees with the three age-related groups observed before for the number of SOX strains across mussel size. Small mussels (≤7cm) and large mussels (14.6cm – 24.1cm) have a low π^SOX^ and harbor one to two strains. Medium-sized mussels (7.2cm – 14.1cm) have a high π^SOX^ and harbor two to nine strains. The community in the largest mussel is an exception, as it has a high π^SOX^, similar to medium-sized mussels, which can be explained by the presence of three strains from two clades.

**Figure 2:**
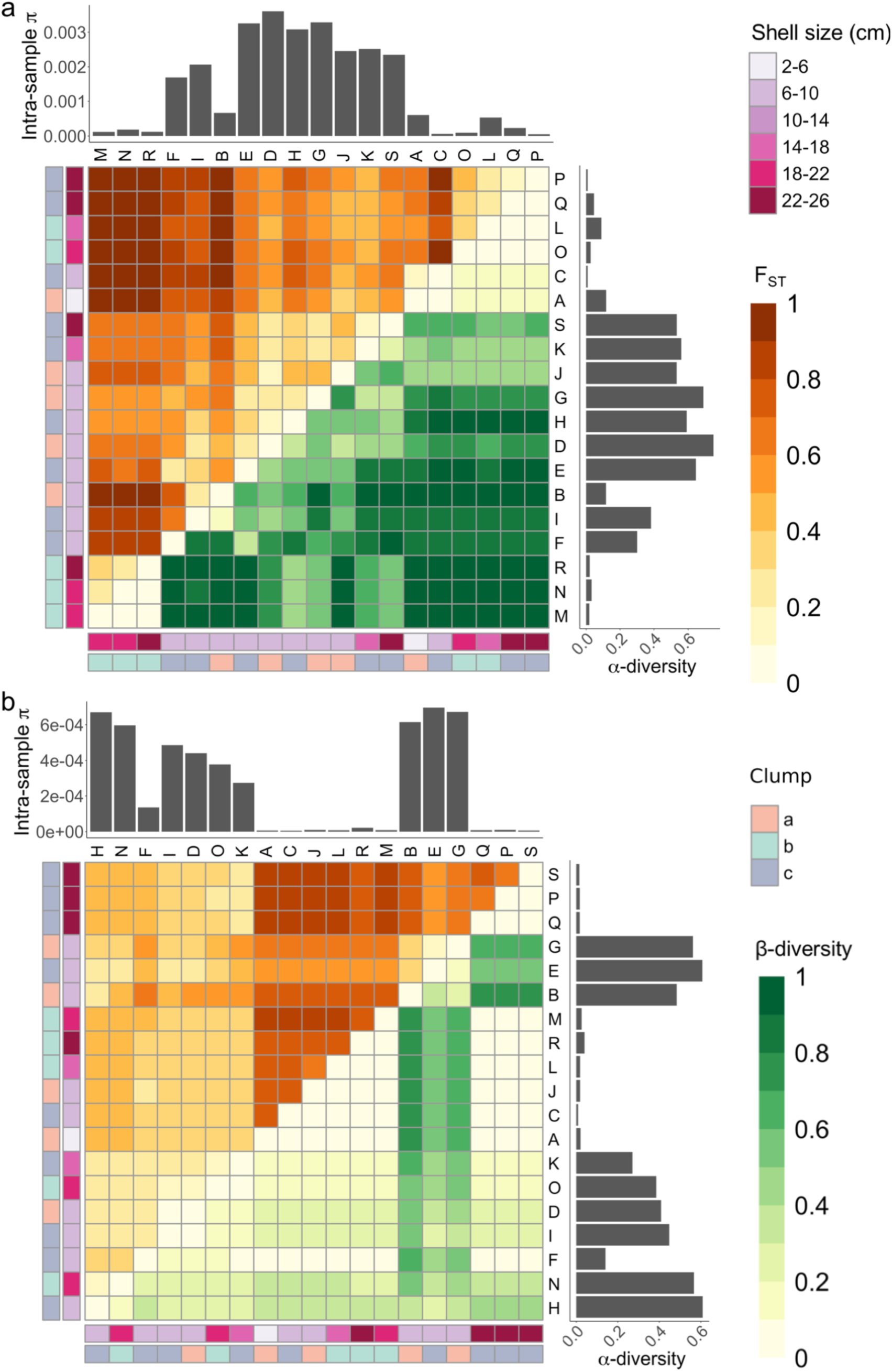
Symbiont population structure for a, SOX and b, MOX. Top left triangle: Intra-sample π and symbiont fixation index (F_ST_) based on SNVs. Lower right triangle: α- and β-diversity based on reconstructed strains. Rows and columns are labelled by sample name, sample location, and shell size. Heatmap hierarchical clustering is based on Euclidean distance of F_ST_. **a**, SOX: mean pairwise F_ST_ is 0.618. Two subpopulations show an extreme degree of isolation: mean pairwise F_ST_ of subpopulation composed of M, N, R, is 0.313; mean pairwise F_ST_ of subpopulation composed of L, O, P, Q is 0.308; mean F_ST_ of sample pairs where one sample is M, N, or R and the other sample is L, O, P, or Q is 0.969. **b**, MOX: mean pairwise F_ST_ is 0.495. The clustering displays two groups: mean pairwise β-diversity of subpopulation composed of A, B, C, D, E, F, G, H, I, J, L is 0.099; mean pairwise β-diversity of subpopulation composed of K, M, N, O, P, Q, R, S is 0.383.

**Table 2.**
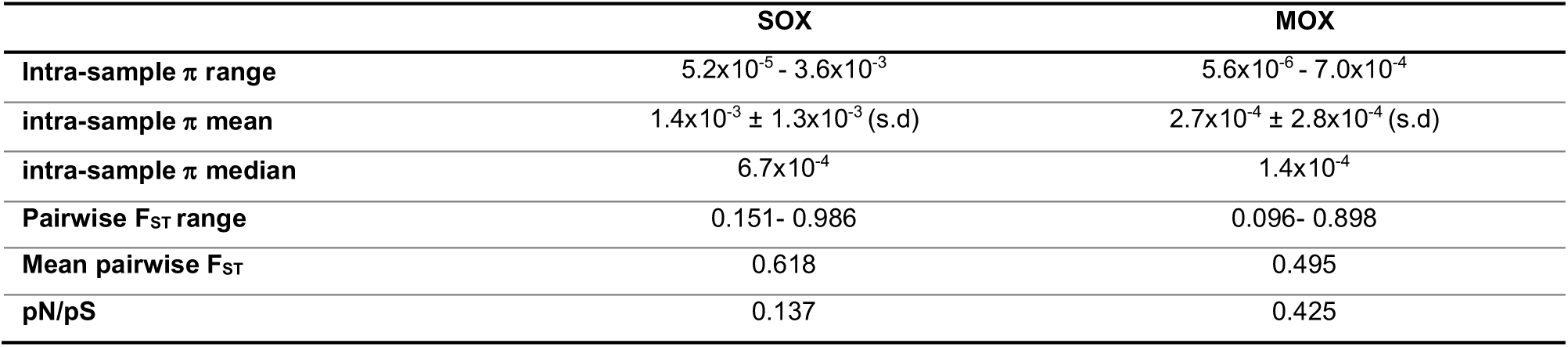
Nucleotide diversity (π), Fixation Index (F_ST_), and pN/pS calculations for both symbiont populations.

|  | SOX | MOX |
| --- | --- | --- |
| Intra-sample $\pi$ range | $5.2 \times 10^{-5}$ - $3.6 \times 10^{-3}$ | $5.6 \times 10^{-6}$ - $7.0 \times 10^{-4}$ |
| intra-sample $\pi$ mean | $1.4 \times 10^{-3} \pm 1.3 \times 10^{-3}$ (s.d) | $2.7 \times 10^{-4} \pm 2.8 \times 10^{-4}$ (s.d) |
| intra-sample $\pi$ median | $6.7 \times 10^{-4}$ | $1.4 \times 10^{-4}$ |
| Pairwise $F_{ST}$ range | 0.151- 0.986 | 0.096- 0.898 |
| Mean pairwise $F_{ST}$ | 0.618 | 0.495 |
| pN/pS | 0.137 | 0.425 |

The MOX nucleotide diversity is significantly lower in comparison to SOX (intra-sample π^MOX^ between 5.6×10^-6^ and 7.0×10^-4^, Table 2, Wilcoxon signed rank test, p=0.015, Fig. 2). Similar to SOX, the MOX α-diversity is significantly positively correlated with π^MOX^ (ρ^2^=0.89, p<10^-6^, Spearman correlation) (Fig. 2b). One group of mussels harbors only MOX strains from clade 2 and is characterized by low MOX nucleotide diversity (A, C, J, L, M, P, Q, R, S, π^MOX^ between 5.6×10^-6^ and 2.1×10^-5^), while the other group habors MOX strains from both clades and is characterized by high MOX nucleotide diversity (B, D, E, F, G, H, I, K, N, O, π^MOX^ between 1.4×10^-4^ and 7.0×10^-4^). These groups are not associated with mussel size. Taken together, we observed a strong correlation between the nucleotide diversity π and α-diversity for both symbionts. Notably, π is based on all the detected SNVs whereas the α-diversity is based only on the strain composition and relatedness. Thus, the strong correlation demonstrates that the strain diversity captures most of the core genome-wide nucleotide diversity.

A comparison of the π values estimated here to other microbiome studies shows that higher π^SOX^ have been observed in other *Bathymodiolus* species (mean between 2.2 ×10^-3^ and 3.9×10^-3^)^25^. The average SOX and MOX nucleotide diversity estimated here is within the range of π values observed in the clam *Solemya velum* microbiome where the symbiont transmission mode is thought to be a mixture of vertical and horizontal transmission^40^. Furthermore, our π estimates are lower than those observed for most bacterial species in the human gut microbiome that are considered horizontally transmitted^19^.

### Geographic isolation of bacterial communities associated with individual mussels

Symbiont transmission mode is an important determinant of the community assembly dynamics^11^. For horizontally transmitted microbiota, similar community composition among hosts may develop depending on factors that affect the community assembly such as the environmental bacterial biodiversity or the order of colonization^41^. To study the degree of geographic isolation between mussel hosts, we calculated genome-wide fixation index F_ST_ and the ecological measure β-diversity at the strain resolution across the metagenomic samples for the two symbionts. Small F_ST_ indicates that the samples stem from the same population whereas large F_ST_ indicates that the samples constitute subpopulations.

Our results revealed generally high pairwise F_ST_ values, indicating a strong genetic isolation between individual mussels (mean pairwise 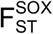 of 0.618, mean pairwise 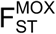 of 0.495, Fig. 2); hence, most mussels in our sample harbor an isolated symbiont subpopulation of SOX and MOX. In addition, the SOX β-diversity is significantly positively correlated with 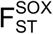 (ρ^2^=0.7, p <10^-6^, Spearman correlation). We observed subpopulations of mussels that are characterized by a low pairwise 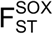 within the subpopulation and a high pairwise 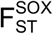 with other mussels. This subpopulation structure is also represented in the distribution of β-diversity (Fig. 2). Thus, mussels from the same subpopulation harbor genetically similar SOX communities and similar strain composition. Examples are one group of mussels including L, O, P, and Q that contains only strains of clade S2 and another group including the mussels M, N, and R that contains only strains of clade S1 (Fig. 2a). Notably, the two subpopulations contain only large mussels that are characterized by a low π^SOX^.

The distribution of pairwise 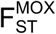 revealed two main groups: one mussel group is characterized by high pairwise 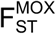 and low π^MOX^ while the other group is characterized by lower 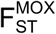 and high π^MOX^ (Fig. 2b). These correspond to the previously described groups, where one contains mussels with a low π^MOX^ and strains from clade M2 and the other group contains mussels with a high π^MOX^ and strains from both clades. We did not observe an association between MOX β-diversity and 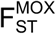 (p>0.05, Spearman correlation), which can be explained by the high proportion of invariant SNVs in MOX. Although the analysis of 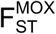 did not reveal MOX subpopulations, the pattern of β-diversity uncovered subpopulations that show a high pairwise 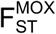. These subpopulations have a low β-diversity and a low nucleotide diversity. One subpopulation consisting of large mussels (P, Q, S) is characterized by the presence of strain M2.3 and the absence of clade M1. Another subpopulation (A, C, J, L, M, R) containing mussels of different sizes is characterized by the dominance of strains M2.1 and M2.2 and the absence of clade M1. Thus, the comparison of strain composition across mussels revealed that the population of MOX is substructured similarly to SOX. However, unlike SOX, the MOX subpopulations are not associated with specific mussel shell sizes.

The high F_ST_ values and the population structure we observed here reveal population stratification, that is especially pronounced for SOX. One possible factor that influences symbiont population structure is host genetics, whose impact on the composition of horizontally transmitted microbiota has been debated in the literature. Studies of the mammal gut microbiome showed that the host genotype had a contribution to the microbiome composition in mice^42^, whereas the association with host genetics was reported to be weak in humans^43^. Analyzing 175 SNVs in 12 mitochondrial genes, we detected no association between mussel F_ST_ and symbiont F_ST_ for any of the two symbionts (Supplementary Information, Supplementary Fig. 8). Consequently, we conclude that the strong subpopulation structure observed for SOX and MOX cannot be explained by mussel relatedness (i.e., host genetics) or location.

Our results provide evidence for a strong genetic isolation between the symbiont populations associated with individual mussels. This finding is consistent with the observed individual-specific symbiont strain composition. In contrast, much lower F_ST_ values were found for SOX populations in other *Bathymodiolus* species sampled from hydrothermal vents (mean F_ST_ per site between 0.05 and 0.17), which implies a weaker genetic isolation in these vents^25^. Our analysis of cold seep *B. brooksi* data revealed SOX subpopulations with low genetic isolation that are observed using both F_ST_, which takes all SNVs into account, and β-diversity at the level of strains. In contrast, only β-diversity disclosed subpopulations for MOX. Thus, strain-resolved metagenomics resolves similarities between individual mussel microbiomes below the species level.

## Discussion

Our analysis revealed strong genetic isolation of symbiotic bacterial populations in individual mussel hosts, indicating geographic isolation between mussels. We hypothesize that this geographic isolation occurs through restricted uptake of SOX and MOX symbionts from the environment over time. The lack of evidence for strong adaptive selection in SOX and MOX strains suggests that the inter-host population structure is the result of neutral processes rather than host discrimination against different strains. Here, we propose a neutral model for symbiont community assembly that explains how restricted symbiont uptake and colonization impose barriers to the symbiont dispersal, which can, over time, lead to inter-host population structure and contribute to the formation of cohesive genetic units within the symbiont population (Fig. 3). In our model, bacteria are acquired from the environmental symbiont pool in juveniles^44^. The presence of a symbiont environmental pool was suggested before based on the detection of symbiont genes in adjacent seawater^45,46^. Nevertheless, the loss of central metabolic enzymes suggests that bacteria disperse in a dormant state^47^. We hypothesize that the dormancy of free-living symbionts and the preservation of few symbiont cells inside bacteriocytes^23^ contribute to the isolation of bacterial populations inside the host cells from the rest of the population, which can lead to recombination barriers. Our results support the self-infection hypothesis^26^, according to which, once the gill is first colonized, bacteria present in ontogenically older tissue infect newly formed gill filaments; thus, the uptake of symbionts from the environment is limited. In addition, decreased growth rate in older mussels may also lead to decreased symbiont uptake. This model provides a plausible explanation for the observed pattern of strong symbiont genetic isolation between mussels and of reduced SOX strain diversity in large mussels. Possible later infections of the gill tissue from the environmental pool may occur due to symbiont loss and replacement driven by environmental changes or increased gill growth rate. Notably, our results are in contrast to a recent study on other *Bathymodiolus* species from hydrothermal vents, that concluded that SOX populations from individual mussels of the same site intermix^25^. This contrast may be explained by differences in the symbiont abundance in the seawater, which is expected to play a role in the colonization process. Our samples originate from a cold seep site with low mussel density (Supplementary Fig. 1); thus, the concentration of symbionts in the surrounding seawater may be correspondingly low. The low symbiont abundance would result in a low probability of later infections and a prevalence of self-infection. In contrast, the symbiont abundance in the seawater at large and densely populated mussel beds at hydrothermal vents is expected to be higher, resulting in a higher probability of later infections.

**Figure 3.**
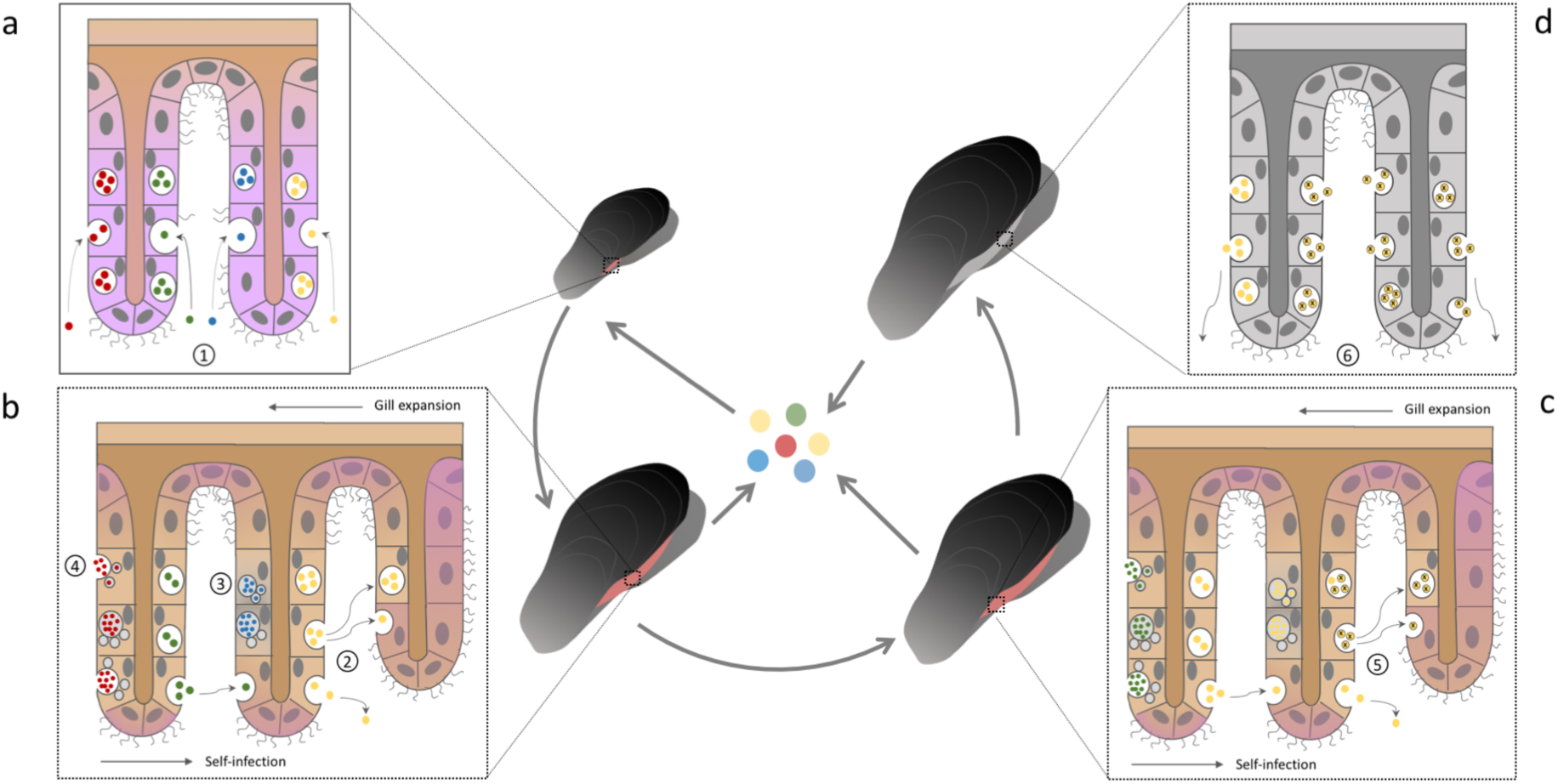
Symbiont colonization dynamics. **a**, The post larvae mussel gill does not take up endosymbionts until the gill presents several filaments and the gill epithelial cells reach a determined developmental stage^26^. At this time point, the filaments are simultaneously infected by different strains via endocytosis (1). This imposes the first bottleneck in the symbiont population, since most likely, not all the strains from the environmental pool can infect the tissue. **b**, Bacteria are released from the host tissue to the environmental pool. As the mussel grows, new filaments are continuously formed in the gill throughout the mussel life span (growing cells shaded in purple). The new tissue is colonized by a self-infection process^26^, which involves infection of the newly formed filaments via endocytosis with bacteria that are released from old tissue via exocytosis (2). The spatial distribution of strains within the gill tissue also supports self-infection^45^. The continuous self-infection process imposes serial founder effects that lead to a reduction in strain diversity, which is mostly driven by drift. Additional sources of diversity loss are: tissue replacement (3) and regulated lysosomal digestion of symbionts^58^ (4). **c**, In older mussels, a unique strain dominates the gill. In addition, *de novo* mutations occur in symbiont genomes (marked by x). Due to serial founder effects within the same mussel, those variants can be quickly fixed inside the mussel (5). **d**, As the mussel dies, bacteria are released from the gill, going back to the environmental pool (6).

The colonization of new filaments over the mussel lifespan via self-infection entails serial founder events on the bacterial population. Throughout this process, new mutations arising in the symbiont population during the lifetime of the mussel can reach fixation due to genetic drift following population bottelnecks. This process is expected to lead to a reduction of symbiont genetic diversity over the mussel life time. Thus, individual mussels develop into independent habitats that harbor individual populations, which are genetically isolated from other mussel-associated symbiont populations and from the environmental pool. The evolution of vertically transmitted endosymbiont populations is similarly affected by serial founder effects^48^, as we suggest here for horizontally transmitted bacteria. However, migration between host-associated populations and the environmental pool results in an increased effective population size for horizontally transmitted bacteria; thus, the population is not subject to the fate of genome degradation as commonly observed in vertically transmitted symbionts^15^. Serial founder effects and recombination barriers due to geographic isolation are important drivers of lineage formation in bacteria^39^. Reduction of genetic diversity due to transmission bottlenecks is considered a hallmark of pathogen genome evolution^49^; examples are *Yersinia pestis*^50^ and *Listeria monocytogenes*^51^. Our model demonstrates that, similar to pathogenic bacteria, genome evolution of bacteria with a symbiotic lifestyle can be affected by serial founder effects due to self-infection.

## Methods

### Collection and sequencing

Twenty-three individuals of *Bathymodiolus brooksi* mussels were collected during a research cruise with the E/V *Nautilus* from the cold seep location GC853 at the northern Gulf of Mexico in May 2015. The mussel distribution at the cold seep was patchy and mussel individuals were collected from three distinct clumps within a radius of 131 meters (coordinates clump a: 28.1237, −89.1404 depth: −1073m, clump b: 28.1241, −89.1401 + depth: −1073m, clump c: 28.1237, −89.1404 + depth: −1073 to 1078m). The gills from each mussel individual were dissected immediately after retrieval and homogenized with sterilized stainless steal beads, 3.2 mm in diameter (biostep, Germany). A subsample of the homogenate for sequencing analyses was preserved in RNA later (Sigma, Germany) and stored at −80°C. DNA was extracted from these subsamples as described by^52^. TruSeq library preparation and sequencing using Illumina HiSeq2500 was performed by the Max Planck Genome Centre in Cologne, Germany, resulting in 250 bp paired-end reads with a median insert size of 400 bp. The raw reads have been deposited in NCBI under BioProject PRJNA508280.

### Construction of the non-redundant gene catalog

Illumina paired-end raw reads from the samples were trimmed for adapters and filtered by quality using BBMap tools^53^. Only reads with more than 30bp and quality above 10 were kept. This results in 37.7 million paired-end reads per sample on average (Supplementary Table 1).

We assembled each of the metagenomic samples individually using metaSPAdes^54^. Genes were predicted *ab initio* on contigs with metaProdigal^55^. These predicted genes were clustered by single-linkage according to sequence similarity using BLAT^56^ (at least 95% of sequence identity in at least 90% of the length of the shortest protein and e-value < 10^-6^). To reduce the potential inflation caused by the single-linkage clustering, we applied two additional filters to discard hits: the maximum ratio allowed between the two compared sequence lengths must be 4 and hits between partial and non-partial genes are discarded. These filters are meant to remove spurious links between sequences due to the presence of commonly spread protein domains. This clustering was performed in two successive steps; first, we obtained sample-specific gene catalogs by performing intra-sample clustering. This is meant to reduce sequence redundancy, resulting in an average of ∼676,000 non-redundant genes per sample (Supplementary Table 1). Second, one-sided similarity search was performed across all pairs of sample catalogs. This resulted in 1,156,207 clusters (26.5%) and 3,207,869 (73.5%) singletons, which make up a catalog of 4,364,076 million non-redundant genes. For each of the clusters, we reconstructed a consensus sequence as cluster representative. To this end, we took the majority nucleotide at each position (ties were resolved randomly).

### Taxonomic annotation of gene catalog

Taxonomic annotation of the gene catalog was performed by aligning the translated genes to the non-redundant protein NCBI database (date: 24/05/18) using diamond^57^ (e-value<10^-3^, sequence identity ≥ 30%) and obtaining the best hit. Genes were annotated as MOX-related if their best hit is *Bathymodiolus platifrons* methanotrophic gill symbiont (NCBI Taxonomy ID 113268) or *Methyloprofundus sedimenti* (NCBI Taxonomy ID 1420851). For SOX, the genomes of thioautotrophic symbionts belonging to four different Bathymodiolus species were used for annotation (NCBI Taxonomy IDs: 2360, 174145, 113267 and 235205). In addition, the gene catalog was screened for mitochondrial genes using best blastp hits against the *Bathymodiolus platifrons* mitochondrial protein sequences (NC_035421.1)^58^ (all e-values <10^-40^). The gene catalog was also screened for symbiont marker genes by best blastp hits to a published protein database for *Bathymodiolus azoricus* symbionts^47^ (80% of protein identity and 100% of query coverage). This allowed to identify 86 SOX and 39 MOX marker genes. The marker gene coverages are generally uniform across a sample, however a high variance in coverage is present in two of the samples (Supplementary Fig. 3). Since the binning method relies on the covariation of coverage across samples, the presence of a high variance in coverage can interfere with the proper clustering of genes, thus, two samples were discarded from further analysis (Dsc1, Dsc2).

### Estimation of the gene catalog coverages

To estimate the gene abundances, we mapped the reads of each metagenomic sample to the gene catalog using bwa mem^59^. Reads below 95% of sequence identity or mapping quality of 20, as well as not primary alignments were discarded. Coverage per position for each gene in the catalog across samples was calculated using samtools depth^60^ and the gene coverage is given by the mean coverage across positions. We first downsampled the reads in each sample to the minimum number of reads found (33M, Supplementary Table 1) and calculated mean coverage per gene to perform the binning and the analyses of coverage variance across symbiont marker genes (see above).

### Genome binning and symbiont core genome identification

Next, we performed co-abundance gene segregation by using a canopy clustering algorithm^29^, which clusters genes into bins that covary in their abundances across the different samples. This approach allows to recover from chimeric associations obtained in the assembly process and to automatically separate core from accessory genes. Gene coverages across samples were used as the abundance profiles for binning. First, genes with a Pearson correlation coefficient (PCC) > 0.9 to the cluster abundance profile were clustered. Then, clusters with PCC > 0.97 between their median abundance profiles were merged and outlier clusters for which the coverage signal originates from less than three samples were removed. In addition, we removed a gene from a cluster if Spearman correlation coefficient to the median canopy coverage profile is lower than 0.7. Finally, overlaps among the clusters were removed by keeping a gene in the largest of the clusters in which it has been found.

This allowed us to cluster 900,310 genes into 98,944 co-abundant gene groups (3 to 699 genes) and three MetaGenomic Species (MGSs, ≥700 genes). An additional filter was applied to the MGSs to obtain final bins by removing outlier genes based on their coverage (Supplementary Fig. 2). To this end, we used the Median Absolute Deviations (MAD) statistic as a cutoff to discard highly or lowly covered genes. We removed genes that are at least 24 times MAD far from the median in at least one of the samples. The bins after outlier gene removal constitute the core genomes of the MGSs. We checked for the completeness of the symbiont bins with CheckM, by screening for the presence of Gammaproteobacteria universal single copy marker genes^61^.

### SNV discovery on the core genomes

To perform single nucleotide variant (SNV) discovery, we mapped the downsampled reads individually for each sample to the gene catalog. Because sample size has been shown to have an effect on variant detection^62^, we normalized the data across samples. To this end, we normalized each sample to the smallest median coverage found in a sample (482x coverage for SOX, 36x coverage for MOX and 568x for mitochdonrial genes). LoFreq was used for probabilistic realignment and variant calling of each sample independently^63^. SNVs detected with LoFreq have been hard filtered using the parameters suggested by GATK best practices^64^. Briefly, SNVs with quality by depth below 2, Fisher’s exact test Phred-scaled probability for strand bias above 60, root mean square of mapping quality below 40, root mean square of base quality above 30, mapping quality rank sum test below −12.5 and read position rank sum test below −8 are kept for further analyses.

The resulting SNVs can be fixed or polymorphic in a sample. Polymorphic SNVs are characterized by the allele frequency of the alternative allele whereas fixed SNVs have an allele frequency of 1. Here, we define SNVs as polymorphic in a metagenomic sample if their frequency is between 0.05 and 0.95 in the sample.

### Population structure analyses

SNV data is used for calculating intra-sample and inter-sample nucleotide diversity (π) as applied before to human gut microbiome species^19^. Intra-sample nucleotide diversity (π) is given as:

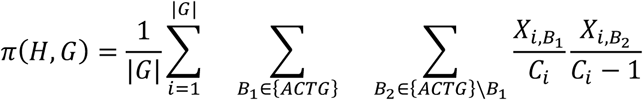

where *H* corresponds to the sample, *G* to the bacterial genome, |*G*| is the length of the analyzed genome and *X*_*i,Bj*_ is the count of a specific nucleotide *B*_*j*_ at a specific locus *i* with coverage *C*_*i*_. Inter-sample nucleotide diversity (π) is then given as follows, where *H*_*1*_ and *H*_*2*_ correspond to the two samples compared:

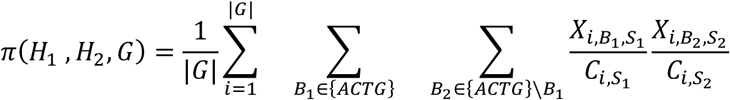

Finally, these diversity measures are used to estimate the fixation index (F_ST_), which measures genetic differentiation based on the nucleotide diversity present within and between populations.

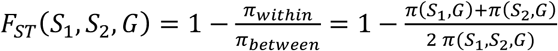

The scripts to calculate genome-wide inter and intra-sample nucleotide diversity (π) and fixation index (F_ST_) across all inter-sample comparisons from pooled SNV data have been deposited at https://github.com/deropi/BathyBrooksiSymbionts.

### Strain deconvolution

We reconstructed the strains for the core genomes with DESMAN^33^. The SNVs with two states and their frequencies in each sample are used by DESMAN to identify strains in the core genomes that are present over multiple samples. Thereby, the program uses the SNV frequency covariation across samples to assign the SNV states to a specific genotype. For SOX, we ran the strain deconvolution five times using different seed numbers and 500 iterations. Due to computational limitations, a subset of 5,000 SNVs was used and the haplotypes considering the whole SNV dataset were inferred *a posteriori*. The five replicates were run for an increasing number of strains from seven to twelve. The program uses posterior mean deviance as a proxy for model fit. A posterior mean deviance lower than 5% was reached in the transition from eleven to twelve strains, therefore the number of inferred SOX strains is eleven. We did not run fewer numbers of strains due to the presence of large posterior mean deviances between runs with a small strain number. Additionally, we ran DESMAN for the SOX dataset that was subsampled to the MOX coverage with no replicates and eleven strains were found using posterior mean deviance. For MOX, we ran four replicates using the whole SNV dataset and 500 iterations. The runs were performed by using an increasing number of strains from two to seven, reaching the optimal number of six strains. The consensus gene sequences of each strain were concatenated to generate the strain core genomes, which were used for further analyses. Splits network of the strain genome sequences were reconstructed using SplitsTree^65^ and uncorrected distances. The position of the root in the splits network was estimated by the minimum ancestral deviation (MAD) method^66^, which uses maximum likelihood phylogenetic trees inferred with IQ-TREE^67^.

### α- and β-diversity

To study the microbial community composition, we estimated α- and β-diversity accounting for strain relatedeness in addition to species richness and eveness. α-diversity was estimated using phylogeny species eveness (PSE)^68^ implemented in the R package ‘Picante’^69^. β-diversity was estimated using the weighted Unifrac distance, which is implemented in the R package ‘GUniFrac’^70^. This measure quantifies differences in strain community composition between two samples and accounts for phylogenetic relationships.

### Allele frequency spectra estimation

The unfolded allele frequency spectra were calculated from biallelic SNVs for each of the bacterial species within individual samples. The unfolded allele frequency spectrum estimation relies on the presence of ancestral states in the population. Because we have no information about the ancestry relationship among the strains present in the samples, we made one main assumption in this regard: the ancestral SNV state in the population corresponds to the one which is present in the higher number of strains. Ties are resolved by arbitrarily assigning one tip of the tree as ancestral state: M2.2 for MOX and S4 for SOX.

### pN/pS and Neutrality Index estimation

We estimated pN/pS for both bacterial populations, which is a variant of dN/dS that can be used based on intra-species SNVs. To this end we first calculated the expected ratio of nonsynonymous and synonymous mutations for each gene by accounting for each possible mutation occurring in each of the codons. Then, we estimated the observed nonsynonymous to synonymous ratio by using the biallelic SNVs. These two measures are later compared, resulting in the pN/pS ratio. pN/pS was estimated genome-wide as well as individually for each of the genes in the two symbiont species. The per-gene pN/pS calculation results into undefinded estimates for genes with no synonymous mutations. To circumvent this limitation, we added 1 to the number of observed synonymous mutations in each gene, which is a standard correction for dN/dS ratios^71^.

The neutrality index (NI) accounts for differences in the ratio of nonsynonymous to synonymous variants between divergent and polymorphic SNVs in order to quantify the departure of a population from neutral evolution^37^. 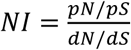, where *pN*, and *pS* are the number of polymorphic synonymous and nonsynonymous sites, respectively, and *dN* and *dS* are the number of divergent synonymous and nonsynonymous sites, respectively. For a coalescent population that evolves neutrally, the nature of fixed mutations that are involved in the divergence of the strains should not be different from that of the polymorphic mutations. An excess of divergent nonsynonymous mutations (NI<1) indicates that the population underwent positive selection or a large demographic change in the past^37^.

Here we used the NI to analyze if differences in selection have been involved in the evolution of SOX and MOX strains. Different strains are typically found in more than one sample and this supports the notion that SNVs that characterize the strains constitute substitutions. We estimated NI by considering two different levels of divergence and polymorphism. First, we defined as divergent all those SNVs that have two possible states among the strains and as polymorphic all the invariant SNVs. Second, we used a more restrictive level of divergence. To this end, we excluded putative recently acquired SNVs from the set of divergent SNVs, by discarding those that have multiple states among strains from the same group. Polymorphic SNVs are all the remaining. The scripts to calculate the allele frequency spectra, pN/pS and NI have been deposited at https://github.com/deropi/BathyBrooksiSymbionts. Statistics and plotting were done in R^72^.

## Acknowledgements

We thank the captain and crew of the E/V *Nautilus* 2015 Expedition, the team of the ROV *Hercules*, the chief scientist on this expedition leg, Eric Cordes, and the expedition leader, Mike Brennan. Additionally, we thank Elie Jami, Robin Koch, Tanita Wein, and Christian Wöhle for critical comments on the manuscript. This work was supported by the CRC1182 *Origin and Function of Metaorganisms*, the Bioinformatics Network at Kiel University, and the North-German Supercomputing Alliance (HLRN, project shb00002). Additional funding support for ND, JMP and RA came from the Max Planck Society, the Gordon and Betty Moore Foundation Marine Microbiology Initiative Investigator Award through grant GBMF3811 to ND and a European Research Council Advanced Grant (BathyBiome, Grant 340535 to ND).

## Author contributions

AK, TD, JMP, and ND designed the study, RA, JMP, and ND collected the data, DRP analyzed the data, DRP, AK, and TD interpreted the results with contribution from RA, DRP, AK, and TD wrote the manuscript with contributions from all authors.

