## Supplement for "Horizontally transmitted symbiont populations in deep-sea mussels are genetically isolated" (from another version of this preprint: version 1, 2019-02-01)

### Supplementary Information

*SOX is the dominant community member in the Bathymodolus microbiota.*

In order to investigate the community composition, we quantified the relative abundance of the microbiota species based on their genomic coverage in the metagenomes. To compare the relative abundances of the two symbionts across the different mussels, we quantified the ratio of SOX to MOX abundance, as well as the symbiont abundance relative to the mitochondrion gene abundance. For this purpose, we annotated twelve *B. brooksi* mitochondrion genes by sequence similarity to the *B. platifrons* mitochondrial genome[1] (Supplementary Fig. 2f). Our results show that SOX has a similar abundance as the mitochondrion, with a median of SOX to mitochondrion ratio = 1.06 (n=19). In contrast, MOX is much less abundant in comparison to the mitochondrion, with a median ratio of 0.0618 (n=19). The two ratios have a low variation across mussel shell sizes (Supplementary Fig. 4a,b). The comparison of SOX and MOX abundance shows that the SOX coverage is on average 19-fold higher than that of the MOX. This shows that SOX is more abundant in the mussel microbiota (Supplementary Fig. 4c). The comparison among sampling clumps shows a higher SOX to MOX ratio in sampling clump c compared to the other two (Supplementary Fig. 4c). This observation is most likely explained by differences in the availability of $H_2S$ and $CH_4$ among clumps, which is a known determinant of SOX and MOX abundance in *Bathymodiolus*[2]. Thus, SOX is the dominant member in metagenomes of the mussel microbiota analysed here, where differences in the SOX to MOX ratio among the mussel metagenomes are likely determined by environmental factors.

*Impact of sequencing depth on diversity analysis*

We tested if the higher SNV density in SOX compared to MOX could be explained by the higher sequencing depth for SOX. To this end, we repeated part of the analyses by analyzing a subset of the orginal SOX reads that were subsampled to achieve MOX coverage levels. However, normalizing SOX to the median MOX coverage of 36x resulted in a SNV density of 12.1 SNVs/kpb. This density is similar to the original estimate of 14 SNVs/kbp and still highly elevated compared to MOX (2.4 SNVs/kbp), indicating that the difference in diversity is not driven by bias due to sequencing depth. Nucleotide diversity of SOX normalized to the MOX coverage (intra-sample $\pi$ between $1.4\times10^{-5}$ and $1.4\times10^{-3}$, mean $4.9\times10^{-4}$, $\pm$ $4.9\times10^{-4}$, s.d.) results in lower estimates than for the full coverage, however, these estimates are still higher than the MOX nucleotide diversity (Table 2).

To test the effect of sequencing coverage on strain inference, we repeated the strain deconvolution for samples where the SOX coverage is decreased to the median MOX coverage. This yielded eleven SOX strains, where some are not identical to the SOX strains reconstructed from the full dataset, yet, the four SOX clades remain well supported (Supplementary Fig. 5). Hence, we consider the strain relationships as more reliable indicators of strain diversity than the number of strains alone. Nevertheless, the resulting AFS does not reveal modes according to the strain clades as observed previously in the full coverage analyses (Supplementary Fig. 9).

*Individual-specific microbiota genetic variants cannot be explained by host genetics.*

Mitochondrial genes have been previously used to identify lineages in *Bathymodiolus* mussels[3]. Here, we used genetic variants in the mitochondrion genome as a marker to investigate the contribution of mussel relatedness to the observed genetic isolation of the symbiont communities. Analysing the twelve mitochondrial genes, we detected 175 SNVs with a density of 15.6 SNVs/kbp (intra-sample $\pi^{Mitochondria}$ between $9.5x10^{-6}$ and $7.7x10^{-5}$, mean $3.3x10^{-5} \pm 2.2x10^{-5}$, s.d.). We found that most of the mitochondrial SNVs are fixed (frequency≥0.95) with an average of 99.8% fixed SNVs per sample. The phylogeny of the dominant mitochondrial haplotype for each sample shows no clustering of individuals according to sample clump or mussel size (Supplementary Fig. 8a), indicating that the mussel individuals analyzed here belong to the same population. The high proportion of fixed SNVs results in high $F_{ST}$ values (Supplementary Fig. 8b,c), as expected for vertically transmitted mitochondrial genomes. We further used the $F_{ST}$ values to detect associations between mussel genetics and symbiont diversity. Our results reveal no association between mussel $F_{ST}$ and symbiont $F_{ST}$ for any of the two symbionts (Supplementary Fig. 8b,c). Consequently, we conclude that the strong subpopulation structure observed for SOX and MOX cannot be explained by mussel relatedness or location.

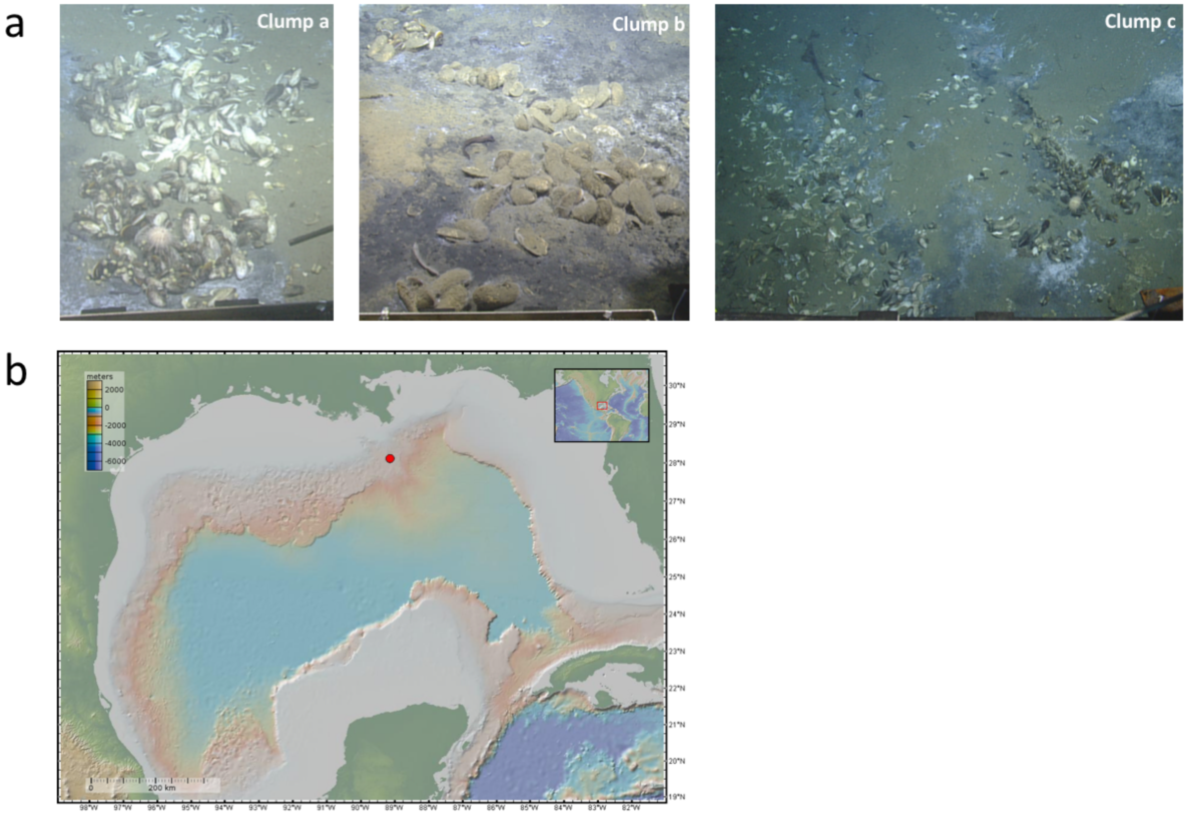


**Figure S1: Sampling information.** a, sampling clumps. b, sampling location at the Gulf of Mexico, different color shades represent sampling depth in meters.

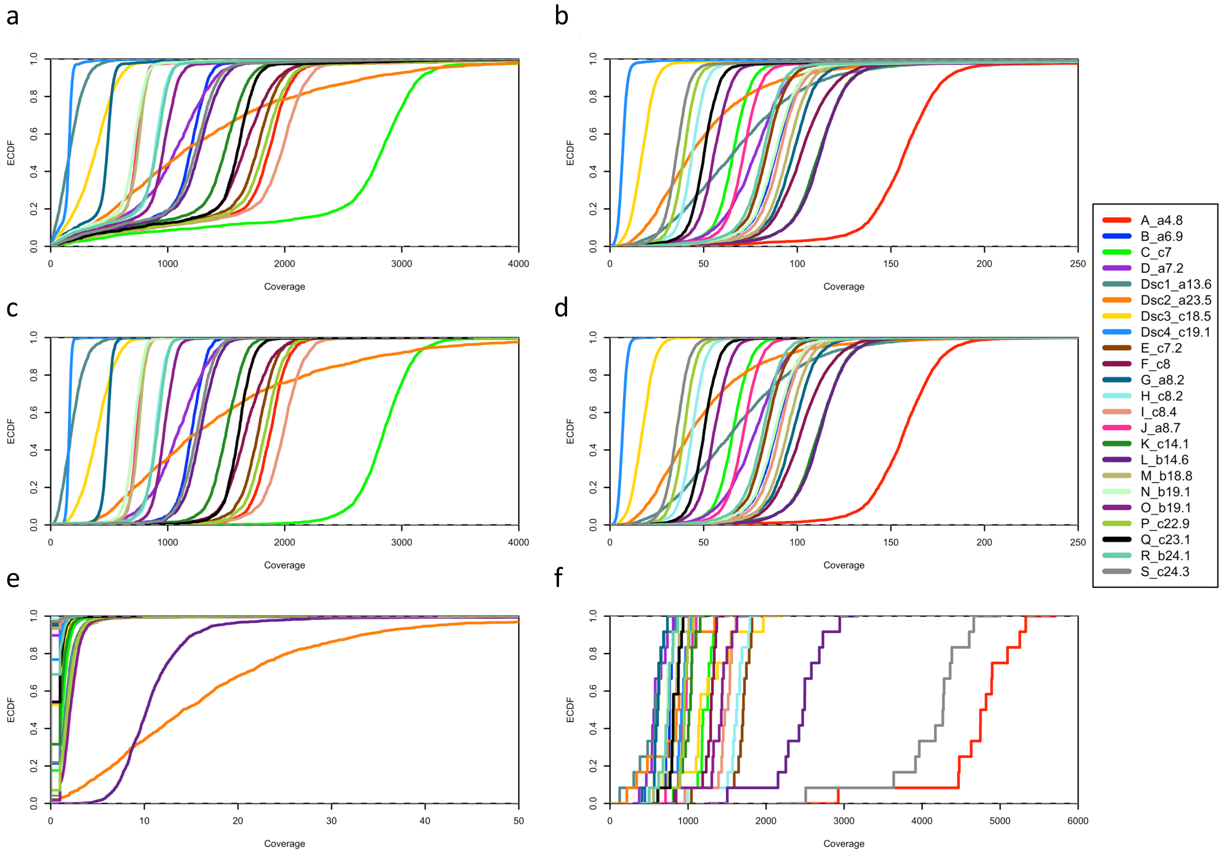


**Figure S2: Coverage of symbiont bins.** Empirical cumulative distribution function (ECDF) of MAGs and mitochondrion. Samples are additionally labeled with the location where they have been sampled and the shell size. 19 samples (A-R) are used for the analysis of population structure and the remaining 4 samples (Dsc1-Dsc4) were discarded during the analysis; Dsc1 and Dsc2 were discarded before the binning due to high variance in symbiont marker gene coverages (Dsc1_a13.6: median coverage=217.41 and SD=98.14 for SOX marker genes, median coverage=70.47 and SD= 41.3 for MOX marker genes; Dsc2_a23.5: median coverage=1368.28 and SD=785.73 for SOX marker genes, median coverage=44.60 and SD=185.12 for MOX marker genes). Dsc3 and Dsc4 were discarded after binning due to low coverage (median coverage < 15x for MOX core genes and median coverage <350x for SOX core genes). Median and standard deviation (SD) across genomes were estimated using only non-discarded samples. **a,** SOX incl. outlier genes (1910 genes, median coverage range: 482-2822, SD range: 157-891), **b,** MOX incl. outlier genes (2618 genes, median coverage range: 36-157, SD range: 97-298), **c,** SOX (1439 genes, median coverage range: 486-2849, SD range: 52-294), **d,** MOX (2518 genes, median coverage range: 36-157, SD range: 6-21), **e,** MGS3 (1,449 genes, median coverage range: 0-10, SD range: 0-11), no outlier genes detected. Total genome length of 1.31 Mbp. The third MGS could not be assigned to a taxonomic level; the top three genera found are Oceanicella (Taxonomy ID:1233054) with 61 genes (4.21%), Neomegalonema (Taxonomy ID 356797, 26 genes, 1.79%), and Micavibrio (Taxonomy ID 213485, 26 genes, 1.79%). This unknown species is present in very low abundance and was mainly found in sample O. **f,** Mitochondrion (median coverage range: 568-4784; SD range: 78-618).

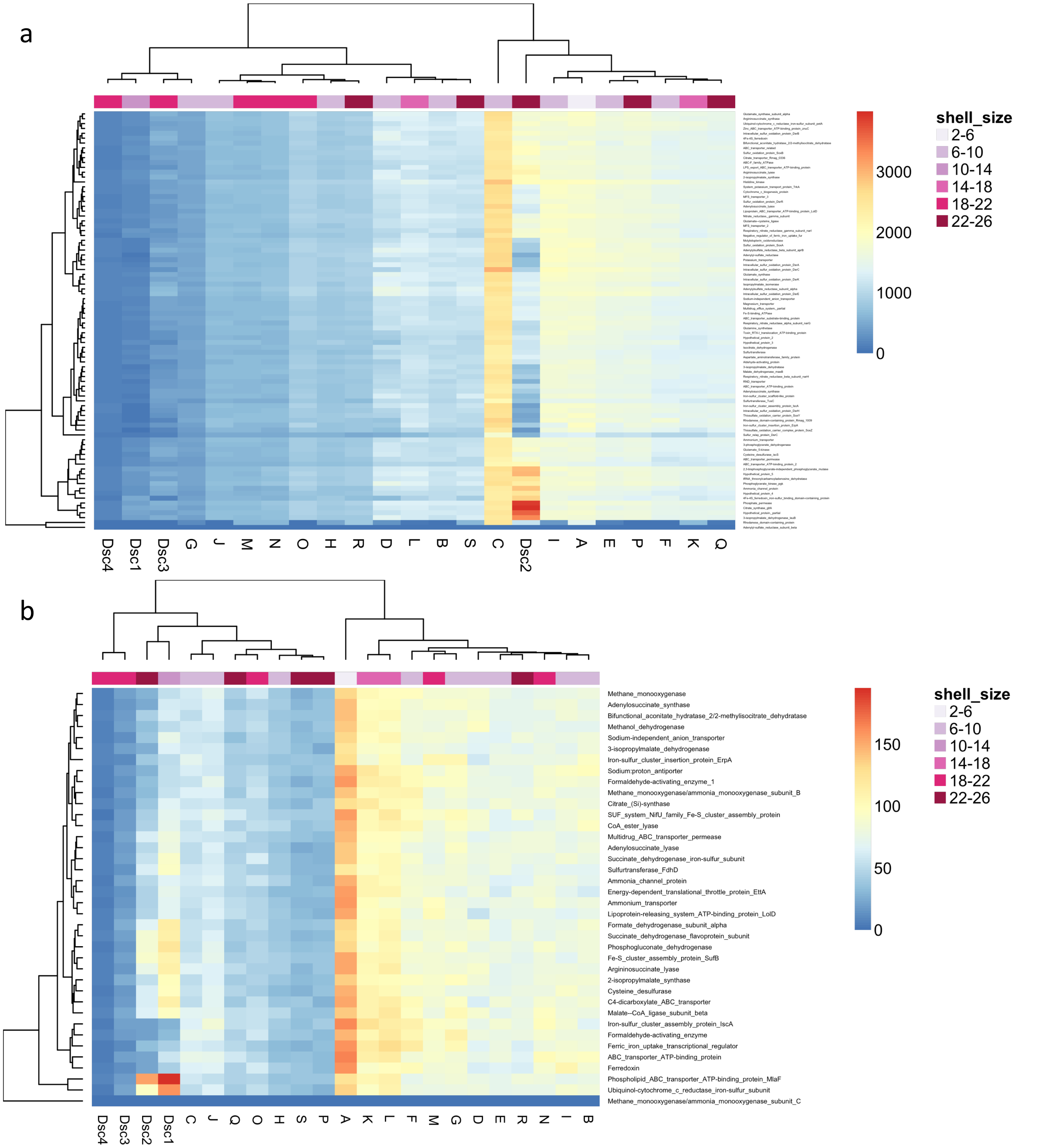


**Figure S3: Gene coverages for a, 86 SOX and b, 39 MOX marker genes.** The two core genomes were validated by screening for the presence of known SOX and MOX marker genes[4]. These genes encode for protein functions in the methane and sulfur/thiosulfate metabolism pathways, such as methane monooxygensase (PmoB), methanol dehydrogenase (XoxF), intracellular sulfur-oxidation proteins (Dsr), sulfur-oxidation proteins (Sox), and adenylsulfate reductase (Apr), as well as for proteins associated to the carbon metabolism, such as malate, succinate and formate dehydrogenases (Mdh, Sdh, and Fdh). A total of 39 MOX and 86 SOX marker genes are present in the gene catalog. Of these, 37 and 77 were included in the MOX and SOX core genomes, respectively.

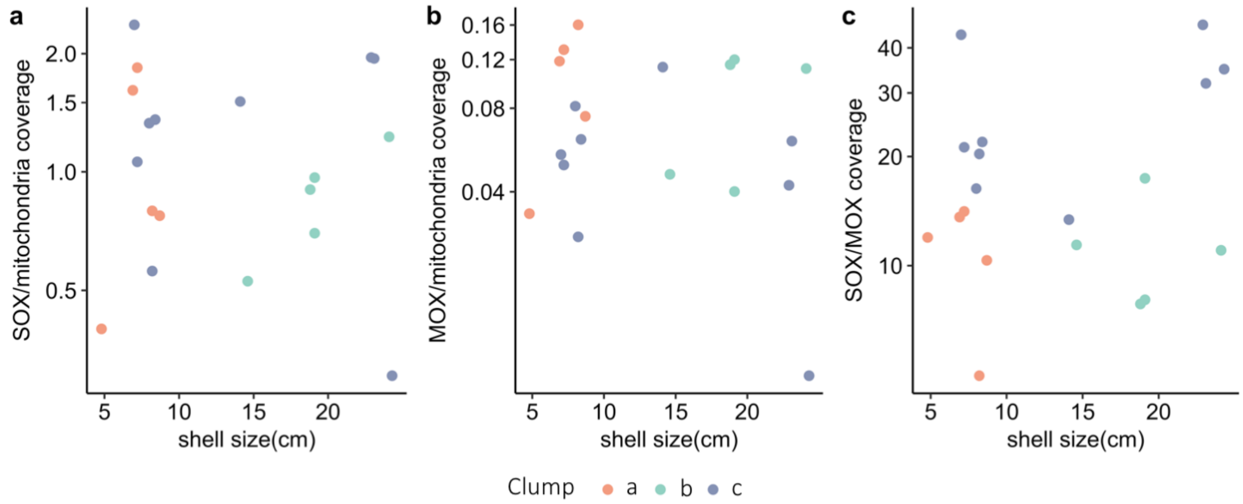


**Figure S4: Relative symbiont abundances across mussels.** Note that the y-axis (ratio) is displayed in log scale. Coverages are estimated by calculating the mean coverage across all genes for SOX, MOX, and mitochondria, respectively.

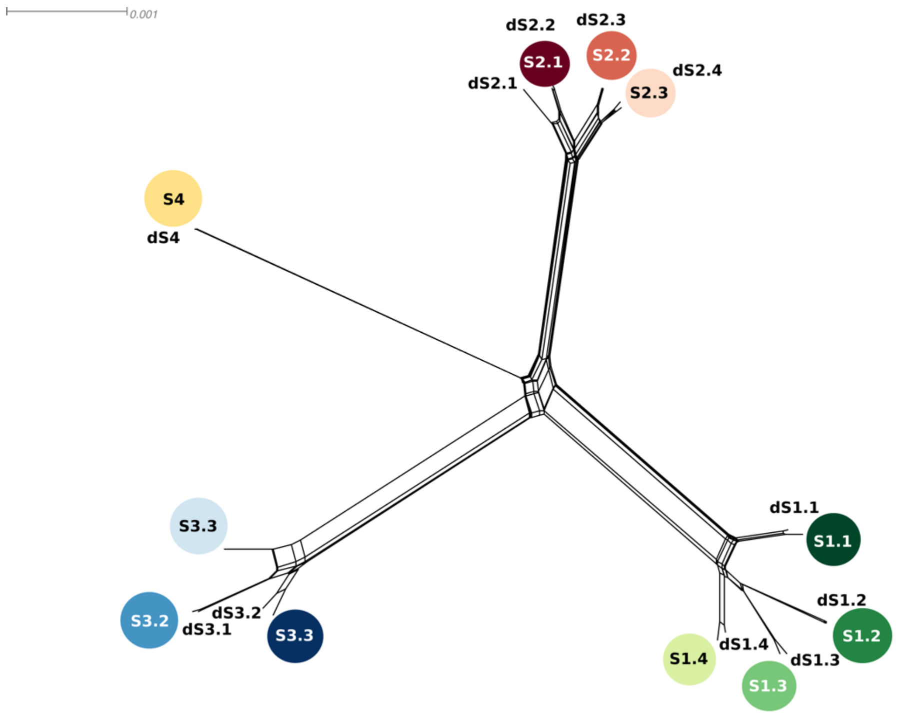


**Figure S5: Splits network for SOX strains reconstructed using the full data set (colored) and SOX strains reconstructed after normalizing SOX to the median MOX coverage (black).**

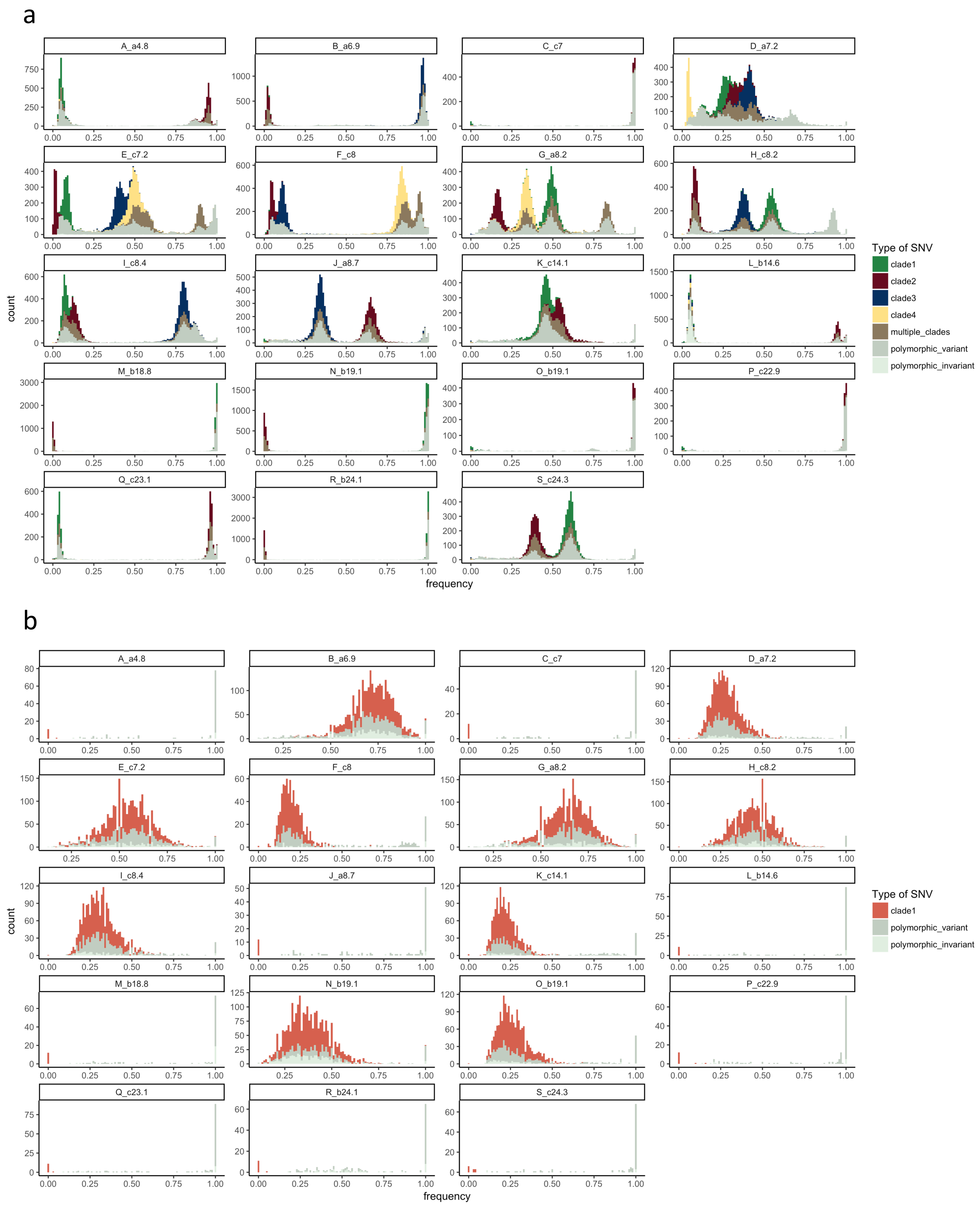


**Figure S6: Intra-sample unfolded allele frequency spectra for a, SOX and b, MOX**. Stacked histogram of the frequencies colored by strain clades (see colors in Fig. 1). SNVs are defined as clade-specific when they occur at the same state in every strain from a clade, as variant, if they differ among strains from one clade and invariant if they are not recovered by the strain deconvolution procedure.

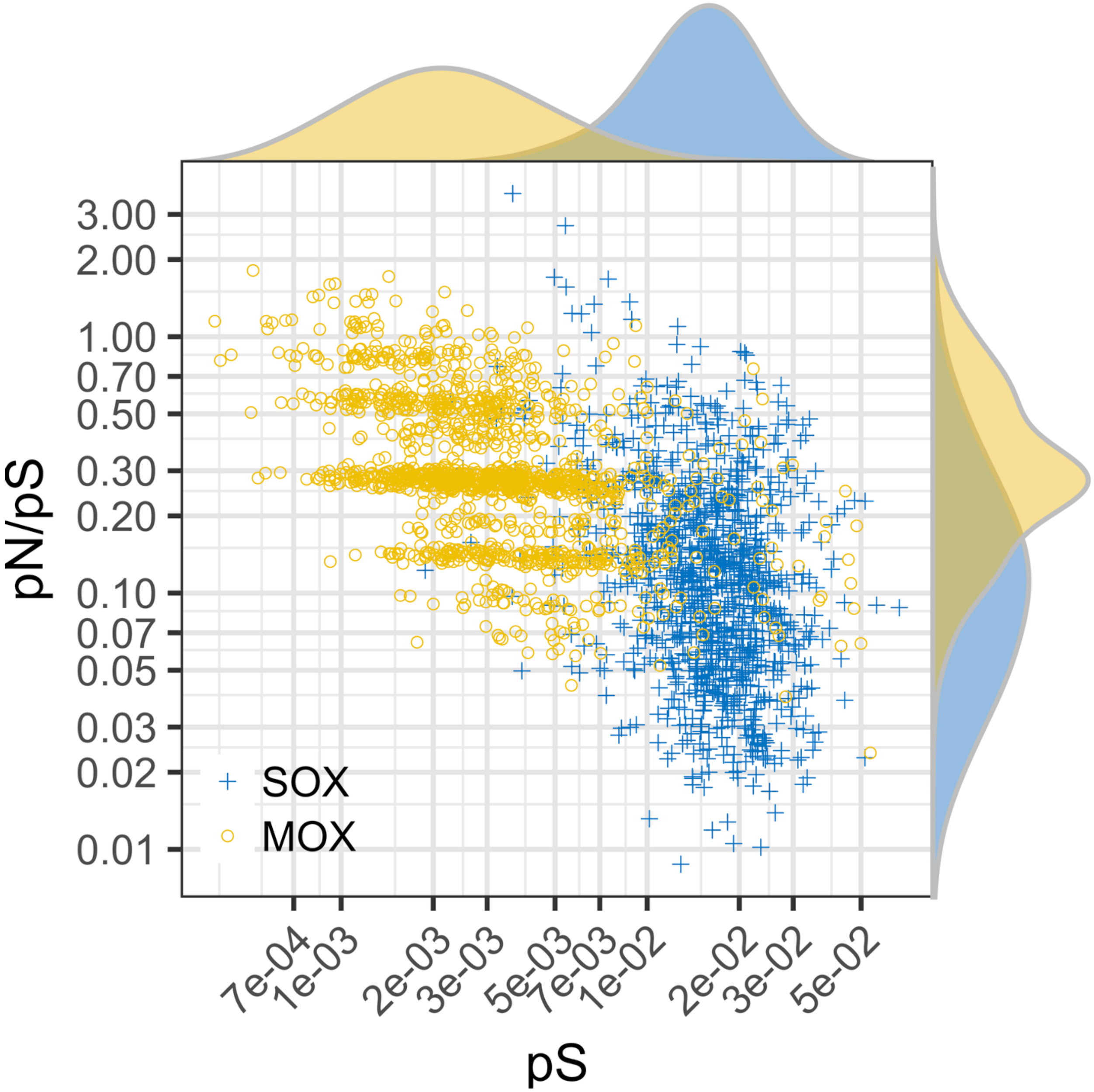


**Figure S7: pS and pN/pS across genes for both symbionts.** Density plots of the distributions are given in the margins. Note the log scale of both axes. Median MOX pS (0.0029) is smaller than median SOX pS (0.015) (Wilcoxon rank sum test, p-value < $10^{-6}$). Median MOX pN/pS (0.28) is larger than median SOX pN/pS (0.12) (Wilcoxon rank sum test, p-value < $10^{-6}$.). Only genes with SNVs are considered (1,117 genes for SOX and 1,359 genes for MOX).

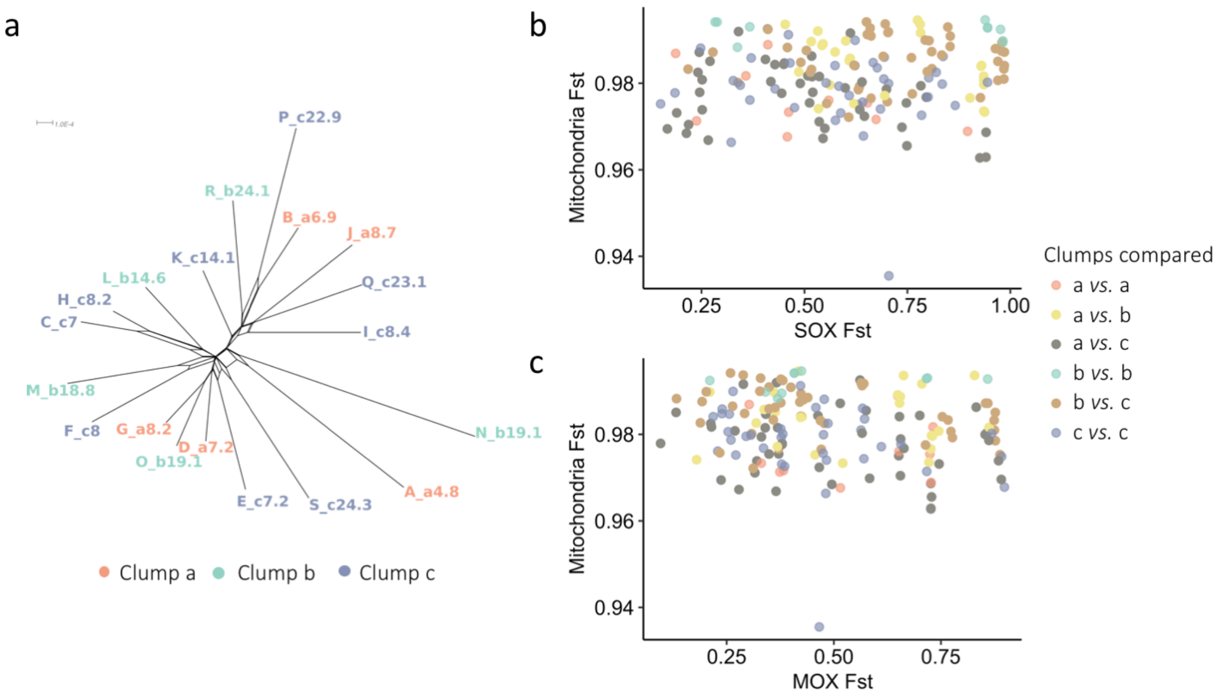


**Figure S8: Mitochondria diversity. a,** splits network[5] (using Uncorrected P distances) based on the dominant haplotypes. We retrieved dominant mitochondrial haplotypes for each of the mussels by identifying SNVs with alternative frequency >0.5 and assigning the alternative nucleotide to these positions. **b,c,** relationship between SOX (b) and MOX (c) $F_{ST}$ and mitochondria $F_{ST}$.

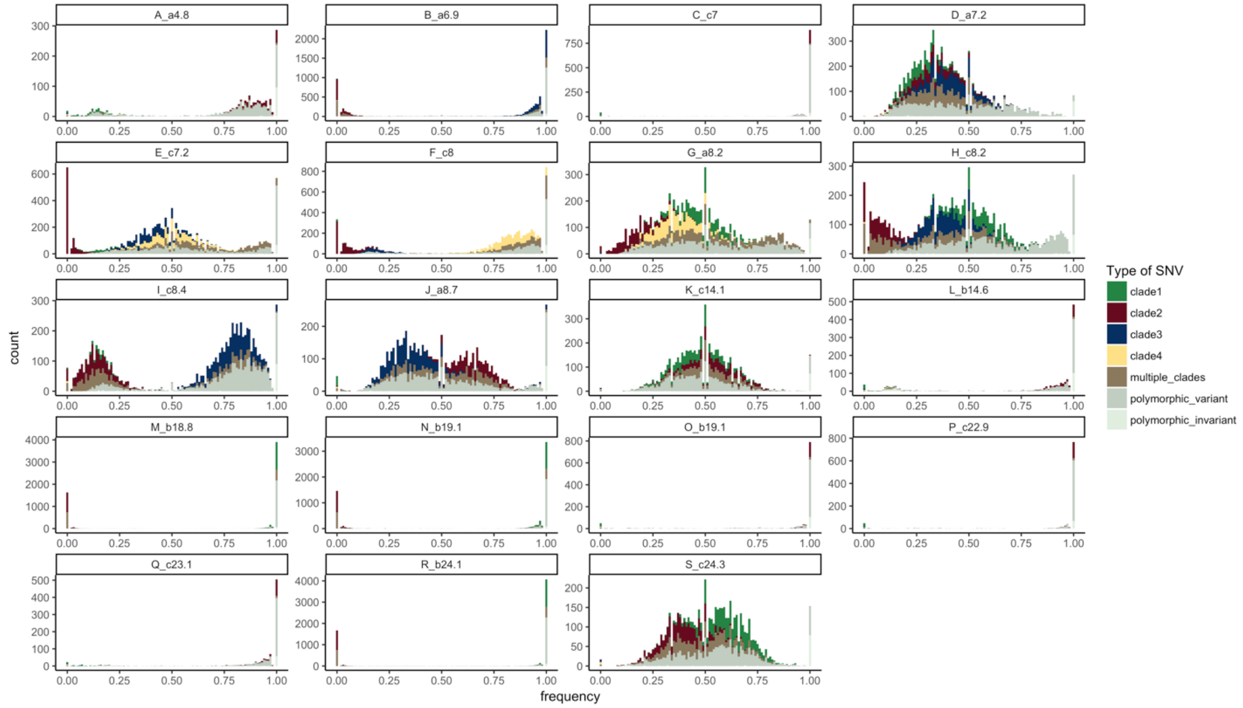


**Figure S9: Intra-sample unfolded allele frequency spectra for downsampled SOX to MOX coverage (~36x)**. See also legend in Supplementary Fig. 6.

**Table S1: Report of raw and processed sequencing data and assembly statistics for the 23 mussel metagenomes from Northern Gulf of Mexico**. # Polymorphic SNVs gives the number of SNVs with a frequency between 0.05 and 0.95. Sample-specific SNVs is the number of SNVs that are present in only one sample and are not detected in any other sample. Dominant strain (≥90%) column shows the strain ID of the dominant strain for these samples containing a dominant strain (strain frequency ≥90%). #Differential dominant strain SNVs gives the number of SNVs that are specific for dominant haplotypes and are retrieved from samples that have a dominant strain (strain frequency ≥90%). We retrieved dominant haplotypes for the mussels with a dominant strain by identifying SNVs with alternative frequency ≥0.9 and assigning the alternative nucleotide to these positions.

| Sample | Size (cm) | Clump | # Reads | # Trimmed reads | # Contigs | # Genes | # Non-redundant genes | # Polymorphic SNVs SOX | # Polymorhic SNVs MOX | # Polymorhic SNVs mitochondria | # Sample-specific SNVs SOX | # Sample-specific SNVs MOX | Dominant strain (≥90%) | #Differential dominant strain SNVs |
|---|---|---|---|---|---|---|---|---|---|---|---|---|---|---|
| A_a4.8 | 4.8 | a | 37,061,240 | 37,028,936 | 1,394,249 | 1,044,714 | 697,354 | 3740 | 30 | 3 | 10 | 6 | - | - |
| B_a6.9 | 6.9 | a | 37,133,252 | 37,106,306 | 1,486,937 | 1,116,948 | 688,707 | 1852 | 3026 | 1 | 140 | 169 | - | - |
| C_c7 | 7 | c | 38,285,976 | 38,252,301 | 1,418,404 | 1,006,084 | 693,453 | 207 | 27 | 0 | 17 | 2 | S2.1 | - |
| D_a7.2 | 7.2 | a | 36,755,366 | 36,731,976 | 1,560,929 | 1,129,326 | 683,199 | 11064 | 2138 | 0 | 20 | 7 | - | - |
| E_c7.2 | 7.2 | c | 37,933,850 | 37,904,916 | 1,498,588 | 1,091,911 | 706,758 | 10505 | 2831 | 1 | 13 | 43 | - | - |
| F_c8 | 8 | c | 38,060,325 | 38,033,705 | 1,432,041 | 1,074,170 | 697,580 | 8923 | 826 | 0 | 242 | 2 | - | - |
| G_a8.2 | 8.2 | a | 37,230,553 | 37,203,081 | 1,453,270 | 1,133,968 | 679,642 | 10404 | 2934 | 0 | 22 | 73 | - | - |
| H_c8.2 | 8.2 | c | 38,070,443 | 38,034,179 | 1,548,988 | 1,150,117 | 699,488 | 10812 | 2717 | 0 | 28 | 51 | - | - |
| I_c8.4 | 8.4 | c | 38,134,036 | 38,103,876 | 1,470,797 | 1,067,621 | 706,089 | 10148 | 2232 | 0 | 52 | 7 | - | - |
| J_a8.7 | 8.7 | a | 37,807,348 | 37,782,857 | 1,463,875 | 1,138,457 | 678,274 | 7147 | 48 | 0 | 49 | 2 | - | - |
| K_c14.1 | 14.1 | c | 36,693,086 | 36,661,232 | 1,460,859 | 1,084,916 | 698,127 | 6742 | 1542 | 0 | 24 | 6 | - | - |
| L_b14.6 | 14.6 | b | 37,412,053 | 37,386,973 | 1,459,725 | 1,106,129 | 687,149 | 3338 | 39 | 0 | 1903 | 3 | - | - |
| M_b18.8 | 18.8 | b | 39,118,344 | 39,086,779 | 1,781,668 | 1,033,575 | 693,534 | 314 | 40 | 0 | 14 | 12 | S1.4 | 42 |
| N_b19.1 | 19.1 | b | 37,259,078 | 37,231,365 | 1,479,982 | 1,141,813 | 691,418 | 397 | 2536 | 0 | 18 | 398 | S1.4 | 74 |
| O_b19.1 | 19.1 | b | 38,172,030 | 38,137,180 | 1,505,470 | 1,132,309 | 707,064 | 315 | 1934 | 0 | 16 | 26 | - | - |
| P_c22.9 | 22.9 | c | 39,623,187 | 39,589,960 | 1,473,410 | 1,101,708 | 693,062 | 162 | 48 | 0 | 10 | 3 | S2.2 | 44 |
| Q_c23.1 | 23.1 | c | 38,172,522 | 38,140,344 | 1,457,910 | 1,097,379 | 683,899 | 615 | 43 | 2 | 12 | 3 | S2.2 | 44 |
| R_b24.1 | 24.1 | b | 37,435,085 | 37,410,384 | 1,492,285 | 1,134,884 | 689,875 | 341 | 89 | 0 | 37 | 44 | S1.4 | 45 |
| S_c24.3 | 24.3 | c | 38,814,311 | 38,775,353 | 1,549,186 | 1,151,312 | 705,170 | 6423 | 32 | 1 | 31 | 5 | - | - |
| Dsc1_a13.6 | 13.6 | a | 33,794,815 | 33,711,379 | 1,599,186 | 1,058,588 | 585,047 | - | - | - | - | - | - | - |
| Dsc2_a23.5 | 23.5 | a | 39,994,621 | 39,945,569 | 1,285,996 | 862,788 | 402,105 | - | - | - | - | - | - | - |
| Dsc3_c18.5 | 18.5 | c | 38,976,397 | 38,944,379 | 1,734,610 | 1,211,676 | 670,826 | - | - | - | - | - | - | - |
| Dsc4_c19.1 | 19.1 | c | 36,555,894 | 36,518,986 | 1,604,292 | 1,176,485 | 703,808 | - | - | - | - | - | - | - |
